# Larval nutrition impacts survival to adulthood, body size, and the allometric scaling of metabolic rate in adult honeybees

**DOI:** 10.1101/2021.02.05.429975

**Authors:** Elizabeth Nicholls, Marta Rossi, Jeremy E. Niven

## Abstract

Resting metabolic rate (RMR) is a fundamental physiological measure linked to numerous aspects of organismal function, including lifespan. Although dietary restriction in insects during larval growth/development affects adult RMR, the impact of larval diet *quality* on adult RMR has not been studied. Using *in vitro* rearing to control larval diet quality, we determined the effect of dietary protein and carbohydrate on honeybee survival-to-adulthood, time-to-eclosion, body mass/size and adult RMR. High carbohydrate larval diets increased survival-to-adulthood and time-to-eclosion compared to both low carbohydrate and high protein diets. Upon emergence, bees reared on the high protein diet were smaller and lighter than those reared on other diets, whilst those raised on the high carbohydrate diet varied more in body mass. Newly emerged adult bees’ reared on the high carbohydrate diet showed a significantly steeper increase in allometric scaling of RMR compared to those reared on other diets. This suggests that diet quality influences survival-to-adulthood, time-to-eclosion, and the allometric scaling of RMR. Given that agricultural intensification and increasing urbanisation have led to a decrease in both forage availability and dietary diversity for bees, our results are critical to improving understanding of the impacts of poor developmental nutrition on bee growth/development and physiology.

**Summary statement:** We show, for the first time, that the nutritional quality of insect larval diets affects the scaling of metabolic rate with body mass in newly emerged adult honeybees.

## 1. Introduction

The resting metabolic rate (RMR) of an organism can account for up to 50% of total energetic expenditure (Morgan, Shelly and Kimsey, 1985) and is intrinsically linked to numerous aspects of physiological and behavioural functioning, from reproductive output to lifespan (Speakman, 2005; Pettersen, Marshall and White, 2018). Despite this, surprisingly little is understood about the drivers of variation in RMR between organisms, particularly at the intra-specific level where consistent individual differences in RMR are frequently observed (McCarthy, 2000; Burton *et al*., 2011). Both across and within many diverse taxa, RMR has been shown to scale allometrically with body size, with larger individuals having higher metabolic rates, and smaller individuals typically having higher mass-specific metabolic rates (Bartholomew, Lighton and Feener, 1988; Gillooly *et al*., 2001; Brown *et al*., 2004; Glazier, 2005; Chown *et al*., 2007). Though the mechanism(s) underpinning allometric scaling of RMR remain highly debated (McNab, 1988; White and Seymour, 2003; Savage *et al*., 2004), scaling exponents have been shown to be affected by several intrinsic and extrinsic factors, including activity level, temperature and diet (Glazier, 2005).

Metabolism is fuelled by food and therefore it is to be expected that an organisms’ diet will have considerable impact on the resources available for energetic expenditure, yet the mechanism(s) by which diet affect RMR and the scaling of RMR remains poorly understood. As highlighted by Naya et al. (2007), in the short term (i.e. hours to days), diet may affect metabolism simply as a result of the energetic processes involved in digestion and absorption of nutrients (Roces and Lighton, 1995; Nespolo, Castañeda and Roff, 2005). In the longer term (i.e. weeks to months), the availability of certain nutrients in an organisms’ diet may affect developmental processes such as organ growth or maintenance processes such as tissue repair (Anderson, 1993; Yang and Joern, 1994). In a number of taxa, including humans, restricting food during developmental stages has been shown to have long-term effects on adult metabolism (Desai and Hales, 1997; Moe *et al*., 2004; Roark and Bjorndal, 2009), potentially allowing organisms to adapt to food scarcity in later life (Hales and Barker, 2001; Wang *et al*., 2016). In many instances, however, organisms are more likely to experience a scarcity of particular nutrients, such as protein or carbohydrates, rather than a complete lack of food, and may be forced to provision their young with sub-optimal, unbalanced diets (Joern, Provin and Behmer, 2012). Yet direct tests of the impact of the nutritional *quality* of developmental diets on adult metabolism are relatively rare outside of epidemiological studies.

Making *a priori* directional predictions about how the nutritional quality of developmental diets might be expected to affect adult metabolic rates is challenging, because theoretical arguments can be made for both positive and negative associations between diet quality and RMR (McNab, 1986; Nussear *et al*., 1998). Nutritional studies have shown that when offered diets of varying composition, organisms defend an optimal intake target of key macronutrients, in particular protein and carbohydrates which provide amino acids and energy vital for survival, growth and reproduction (Karasov, Martínez Del Rio and Caviedes-Vidal, 2011; Simpson and Raubenheimer, 2012; Roeder and Behmer, 2014). Optimal intake targets can be achieved through behaviours such as selective or compensatory feeding, or physiological/morphological means such as increasing gut length or food retention time (Felton, 1996; Behmer, 2009; Burton *et al*., 2011). Though insects have long been used as models to study the regulation of nutritional intake targets (Behmer, 2009) studies of the long-term impact of variation in nutrition over the course of development are somewhat lacking (Roeder and Behmer, 2014), and studies of the subsequent effects on adult metabolism are largely non-existent. A recent study found that adult stick insects exhibit developmental diet dependent differences in RMR when reared from birth on leaves from plant species varying in their nutritional content and digestibility (Hill, Silcocks and Andrew, 2020), but the impact of developmental diet on the scaling of RMR and body mass was not considered. Shorter term studies conducted in adult insects only are more common, and have typically observed a reduction in RMR in response to a lower quality diet (Zanotto *et al*., 1997; Ayayee *et al*., 2018, 2020, but see Clark, Zera and Behmer, 2016).

Bees meet all their nutritional demands *via* pollen and nectar collected from flowers (their main source of protein and carbohydrate respectively), and unlike the nymphs and larvae of traditional models of insect nutrient regulation, such as locusts and caterpillars, bee larvae are entirely dependent on the provisioning choices of adult bees. This means bee larvae likely have very little opportunity to selectively regulate their intake of nutrients (but see Austin and Gilbert, 2018). Honeybees are unique among bees in that in-hive nurse bees process the pollen and nectar brought back to the nest by foragers, and convert it to a nutritional substance known as royal jelly which they then regurgitate for larvae (Wright, Nicolson and Shafir, 2018). Containing approximately 60% water, 15% protein, 20% carbohydrates, and 5% fats, the exact macronutrient content of royal jelly can vary between colonies and over time (Howe *et al*., 1985; Garcia-Amoedo and De Almeida-Muradian, 2007; Ferioli, Armaforte and Caboni, 2014). Furthermore, a recent study has demonstrated that nurse honeybees are unable to discriminate between pollen diets on the basis of nutritional quality (protein and/or lipid content) (Corby-Harris *et al*., 2018), meaning the proportion of macronutrients that individual larvae receive in their diet could vary, particularly in times or areas where the diversity of forage is limited (Donkersley *et al*., 2017). In addition, there is recent evidence to suggest that rising CO_2_ levels associated with climate change are negatively affecting the nutritional quality of pollen and nectar provided by plants (Ziska *et al*., 2016). Given widespread concerns regarding the combined effects of habitat degradation and agricultural intensification on the availability of sufficiently diverse floral resources to meet the nutritional needs of adult bees and their offspring (Naug, 2009; Brodschneider and Crailsheim, 2010; Donkersley *et al*., 2017), and the fact that bees provide a pollination service vital to global food security, the question of how developmental diets impact on the metabolic function of adult bees is extremely apposite.

Here we used *in vitro* rearing methods to tightly control honeybee larval diets independent of nurse bee behaviour, permitting an examination of the impact of diet nutritional composition on honeybee development and adult physiological function. Previous studies have shown that the ratio of protein to carbohydrate in honeybee larval diets can have significant impacts on larval survival (Helm *et al*., 2017), with unbalanced diets heavily skewed to either macronutrient resulting in poor growth and survival. To our knowledge this is the first study to test the RMR of adult bees reared on different larval diets *in vitro*. By manipulating the ratio of royal jelly (protein) to sugars (carbohydrates), we aimed to determine the impact of specific macro-nutrients on adult RMR and scaling with body size.

## 2. Materials and Methods

Honeybee *(Apis mellifera* L.) larvae were obtained from full-sized colonies housed on the University of Sussex campus, and reared in the laboratory using the *in vitro* method described by Schmehl *et al*. (2016). Briefly, three-day old larvae were removed from the comb using a grafting tool, transferred to individual wells of a 48-well cell culture plate, and placed into an incubator fixed at 35°C, 94% relative humidity (RH). Larvae were fed once per day for five days, and upon pupation transferred to a fresh cell culture plate. Survival was monitored daily until adult emergence.

### Diet manipulation

A standard *in vitro* rearing diet (Table 1) of yeast (Sigma-Aldrich UK), royal jelly (The Raw Honey Shop, Brighton) and sugars (glucose and fructose, Sigma-Aldrich UK) was manipulated to contain differing amounts of protein (using royal jelly as a proxy) and/or carbohydrate (glucose and fructose), following the methods of Helm *et al*. (2017). Larvae were reared on one of five diets (Table 1, D1-5), where the amount of protein and carbohydrate was either increased or decreased relative to the diet described by Schmehl *et al*. (2016).’ Royal jelly was stored frozen at −20°C in 50 mL aliquots. Diets were freshly made every two days and stored at 4°C. Larvae were fed once per day for five days, and the volume of food varied according to the day of the experiment (Days 1 and 2 = 10 μL; Day 3 = 20 μL; Day 4 = 30 μL; Day 5 = 40 μL; and Day 6 = 50 μL). Between 60 and 78 larvae were assigned to each treatment group *(N* = 371 larvae in total; D1=78; D2=60; D3=78, D4=78; D5=77). Bees were reared in two cohorts, grafted on 30/9/2019 and 20/10/2019. Royal jelly nutritional values (supplementary data) were obtained by the supplier (The Raw Honey Shop, Brighton) using the international standard for royal jelly (ISO 12824:2016). From these values we calculated the proportion and ratio of protein:carbohydrate (P:C) in each of the five diets (Table 1).

**Table 1.** Composition of artificial diets fed to larvae in the P:C ratio diet manipulation experiment (P = Protein, C = Carbohydrate).

| **diet** | **P** | **C** | ***n*** | **% diet component** |  |  |  |  |  |  | **P:C**<br>**ratio** |
| --- | --- | --- | --- | --- | --- | --- | --- | --- | --- | --- | --- |
|  |  |  |  | yeast | royal<br>jelly | glucose | fructose | water | total<br>P | total<br>C |  |
| **D1** | med | med | 78 | 1.0 | 51.0 | 4.1 | 8.2 | 35.7 | 8.2 | 18.5 | 1:2.3 |
| **D2** | med | high | 60 | 1.0 | 48.1 | 5.8 | 11.5 | 33.7 | 7.7 | 23.2 | 1:3.0 |
| **D3** | med | low | 78 | 1.1 | 54.3 | 2.2 | 4.3 | 38.0 | 8.7 | 13.1 | 1:1.5 |
| **D4** | high | med | 78 | 0.9 | 57.5 | 3.5 | 7.1 | 31.0 | 9.2 | 17.6 | 1:1.9 |
| **D5** | low | med | 77 | 1.2 | 42.2 | 4.8 | 9.6 | 42.2 | 6.8 | 19.5 | 1:2.9 |

### Measuring resting metabolic rates

To determine how larval diet affects adult metabolism, the RMR of adult bees was measured on the day of emergence (between 14-17 days from the day of grafting) using flow-through respirometry, with CO_2_ production used as a measure of metabolic rate. Emerging adults were first individually weighed to the nearest mg using a precision balance (Mettler Toledo, UK). Bees were then restrained using a small cylinder of metal mesh to allow gas exchange, before being placed into a 2 mL plastic chamber. Air scrubbed of CO_2_ and H_2_O was then pumped through the chamber at a consistent rate of 100 mL min^-1^ via a mass flow controller (GFC17; Aalborg, NY, USA), before passing through an infrared CO_2_-H_2_O analyser (Li7000, Li-Cor) which captured data on CO_2_ production, relative to an empty control chamber (Nicholls *et al*., 2017; Perl and Niven, 2018). The temperature in the room was held constant at 25°C (± 2°C) and recordings lasted for 20 minutes per bee. The first five minutes of the recording were treated as a settling period for the bee to adjust to the experimental set up and were excluded from analysis. During recording the plastic chamber was covered to ensure it was dark, which reduced bee movement. The order in which bees from different diet treatment groups were measured was randomised. After recording, bees were frozen to immobilise them, and digital callipers were used to measure the intertegular span (defined as the distance between the points at which the wings attach to the thorax) in mm, a proxy measure for body size (Cane, 1987).

### Data analysis

Respirometry data was analysed using OriginPro software (Origin 2016, OriginLab Corporation, Northampton, MA, USA). Volumes of CO_2_ were baseline corrected and temperature normalised using the Q10 correction for temperature differences. To calculate the rate of CO_2_ production per bee, the volume of CO_2_ (ppm) was converted to CO_2_ fraction and multiplied by the flow rate (100 mL min^-1^). The integral of CO_2_ min^-1^ *versus* min was calculated for a stable 15-minute period of the recording, and divided by this time to give a rate of μl CO_2_ h^-1^.

All statistical analyses were conducted in R 3.6.2 (R Core Team, 2019. https://www.R-project.org). To examine how diet quality impacts larval survival, Kaplan-Meier survival analysis was performed using the survfit function from the ‘survival’ package. The log-rank test was used to test for differences in survival between diet treatments with a Bonferroni correction for multiple comparisons. Linear and mixed effect models were performed by restricted maximum likelihood (REML) estimation using the lmer and glmer function from the ‘lme4’ package to test the impact of diet treatment on the time to adult emergence (days), wet body mass (mg), body size (using intertegular distance as a proxy measure; mm), body condition (body mass/body size; mg/mm) and CO_2_ production (μL CO_2_ h^-1^). The continuous variables body mass, body size, body condition and CO_2_ production were log transformed. Date of grafting was included as a random effect. For all models, Diet 2 was used as the reference category because bees in this treatment had the best survival. Significances of the fixed effects were determined using Satterthwaite’s method for estimation of degrees of freedom by using the anova function from ‘lmerTest’. Estimated marginal means (emm) and pairwise comparisons were obtained using the ‘lsmeans’ package and the *p*-value adjusted with the Tukey method. To test for differences in variance, we used the Brown-Forsythe test for non-normal data. All plots were made using the ‘ggplot2’ package.

## 3. Results

### The ratio of P:C in larval diets affects honeybee development and survival

Diet had a significant effect on the survival of honeybees to adult emergence (Fig. 1, Table S1,2; Kaplan-Meier log-rank test, *X*^2^_4_ = 54.7, *p* < 0.001). Larvae reared on the high carbohydrate diet (D2), which had a P:C ratio of 1:3, had the best survival (70%), significantly higher than all other treatment groups (Fig. 1, Table S2; Log-rank test D2-D1p=0.023; D2-D3 p <0.001; D2-D4 p <0.001; D2-D5 p=0.013)). Bees reared on the high protein diet (D4, P:C 1:1.9) had very poor survival (22%), and only one bee reared on the low carbohydrate diet (D3, P:C 1:1.5) survived to adulthood (Fig. 1). Consequently, bees from D3 are excluded from subsequent analyses. Bees reared on the diet recommended by Schmehl *et al*. (2016) for rearing larvae (D1, P:C 1:2.3), and the low protein diet (D5, P:C 1:2.9) had similar levels of survival (Table S1,2), with just under half of all larvae reaching adulthood (~45%, Fig. 1). Diet also had a significant effect on development time (days to emergence) (Table 2, Table S3; *X*^2^_3_ = 22.14, *p* < 0.001), with bees reared on the high carbohydrate diet that maximised survival (D2) taking significantly longer to emerge (emm ± s.e. = 16.0 ± 0.96 days) than those in all other treatment groups (D1 = 15.5 ± 0.96; D4 = 15.3 ± 0.97; D5= 15.7 ± 0.96 days).

**Figure 1.**
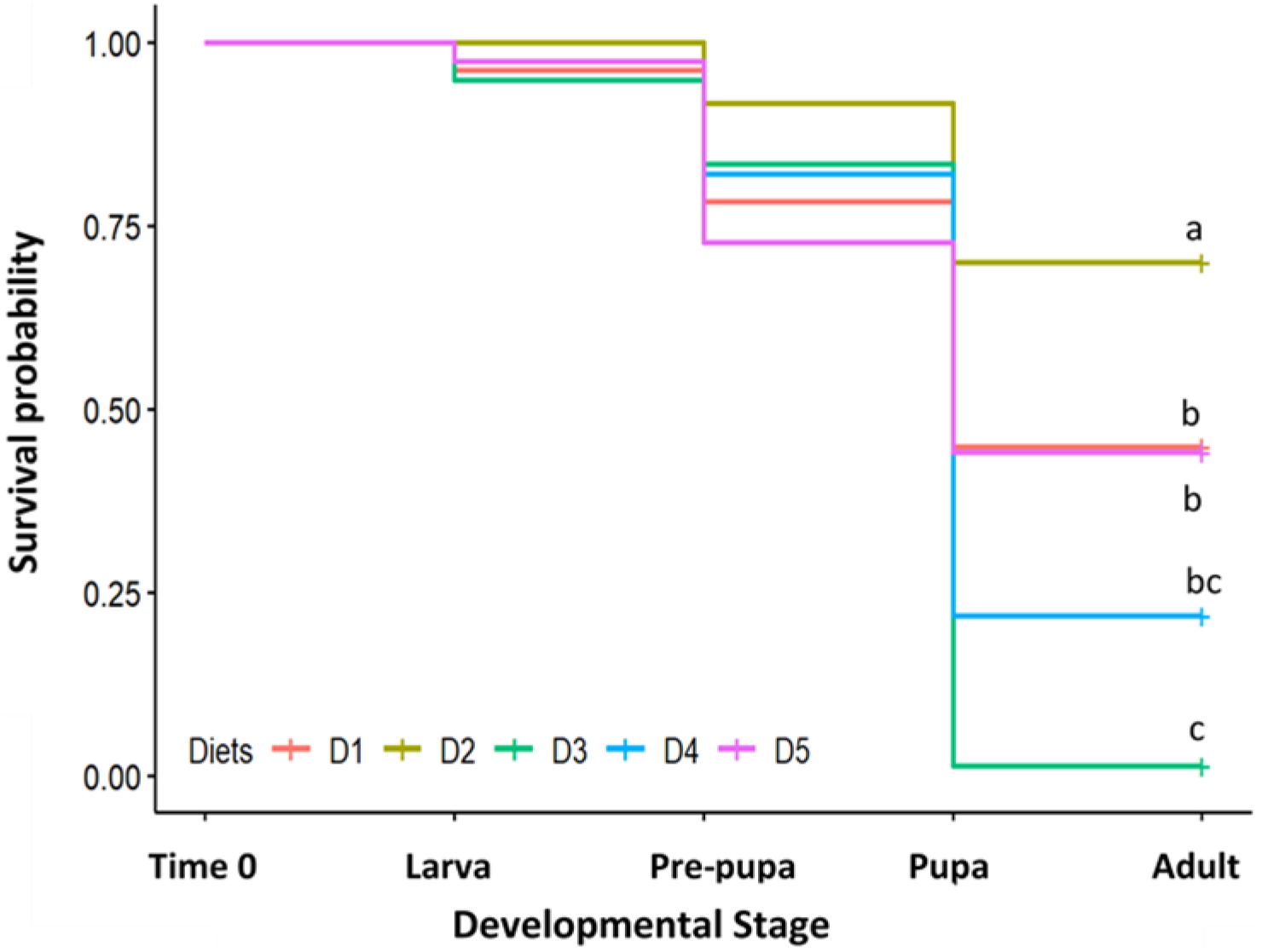
Probability of survival of bees reared on different larval diets (P:C ratio in diets: D1 = 1:2.3; D2 =1:3.0; D3 = 1:1.5; D4 = 1:1.9; D5 = 1:2.9). The number of larvae in each treatment group at Time 0 is as follows: D1= 78; D2=60; D3=78; D4=78; D5=77. Crosses indicate the proportion of individuals in each diet treatment that reached adulthood (censored data). Letters indicate statistically significant differences in survival *(p* < 0.05, Kaplan-Meier analysis).

**Table 2.** Least-square pairwise comparisons of the effect of diet treatment on the time to adult emergence, body mass on emergence, body size and body condition. Models applied were (days to emergence ~ diet + (1|grafting cohort)), (body mass ~ diet + (1|grafting cohort)), (body size ~ diet + (1|grafting cohort)) and (body condition ~ diet) respectively. *P*-values were adjusted using the Tukey method. (P:C ratio in diets: D1 = 1:2.3; D2 =1:3.0; D4 = 1:1.9; D5 = 1:2.9). The number of bees measured in each treatment is as follows: D1=28; D2=33; D3=0; D4=10; D5=30. See Table S3 for the complete outcome of the models.

| diet | time to emergence<br>(days) |  | body mass<br>(mg) |  | body size<br>(mm) |  | body condition<br>(mg/mm) |  |
| --- | --- | --- | --- | --- | --- | --- | --- | --- |
| | estimate ± s.e. | *p*-value | estimate ± s.e. | *p*-value | estimate ± s.e. | *p*-value | estimate ± s.e. | *p*-value |
| D1 – D2 | **-0.50 ± 0.13** | **<0.001** | 0.91 ± 3.07 | 0.991 | -0.00 ± 0.05 | 1.000 | **3.40 ± 1.14** | **0.021** |
| D4 – D2 | **-0.63 ± 0.18** | **<0.001** | -9.23 ± 4.35 | 0.154 | **-0.24 ± 0.06** | **0.002** | 2.40 ± 1.38 | 0.315 |
| D5 – D2 | **-0.36 ± 0.13** | **<0.025** | 0.68 ± 4.22 | 0.996 | -0.07 ± 0.05 | 0.500 | 2.70 ± 1.13 | 0.890 |
| D4 – D1 | -0.18 ± 0.17 | 0.714 | -10.14 ± 3.07 | 0.083 | **-0.24 ± 0.05** | **<0.001** | -1.00 ± 1.40 | 0.090 |
| D5 – D1 | 0.14 ± 0.12 | 0.676 | -0.23 ± 2.99 | 0.100 | -0.07 ± 0.04 | 0.333 | -0.70 ± 1.14 | 0.927 |
| D5 – D4 | 0.32 ± 0.17 | 0.243 | 9.09 ± 4.17 | 0.088 | **0.17 ± 0.05** | **0.008** | 0.30 ± 1.38 | 0.996 |

### The ratio of P:C in larval diets affects adult body mass, size and condition

On emergence, bees reared on the high protein diet (D4), the second worst diet for survival, weighed approximately 10 mg less on average than those reared on all other diets (Fig. 2A), and were significantly lighter than those reared on the high carbohydrate diet D2 (Table 2, Table S3; estimate ± s.e. −9.23 ± 4.32 mg, df = 96.61, *p* = 0.035). Variance in body size also differed between diet treatments (Fig. 2A). There was a significant difference in the variance of body mass, both between bees reared on D2 and D1 (Table S4; Brown-Forsythe Test, *p* = 0.007), and D2 and D5 (Brown-Forsythe Test, *p* = 0.016) suggesting that the diet maximising survival (D2) allowed for a greater range of body masses.

**Figure 2.**
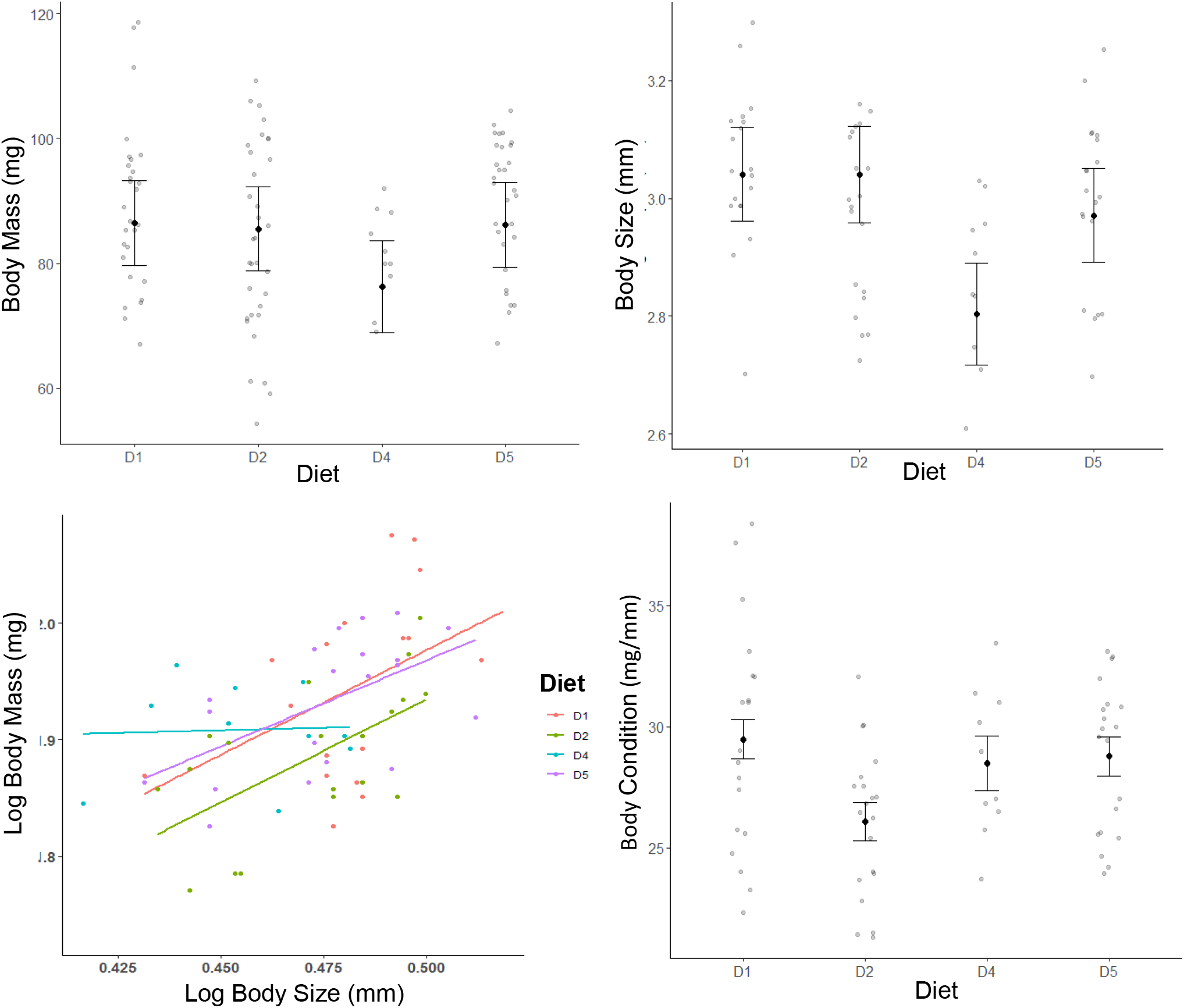
Body mass (A) body size (B) the scaling of body mass and body size (C) and body condition (body mass/body size) (D) of adult bees in each diet treatment. Grey points are the individual data points, black points represent the estimated marginal mean and whiskers are the standard error of the mean. (P:C ratio in diets: D1= 1:2.3; D2 = 1:3.0; D4 = 1:1.9; D5 = 1:2.9). The number of bees measured in each treatment is as follows: D1=28; D2=33; D3=0; D4=10; D5=30.

Bees reared on the high protein diet (D4) were also significantly smaller (emm ± s.e. = 2.80 ± 0.09 mm) than bees in all other treatment groups, as measured by the intertegular span (Fig. 2B; Table 2, Table S3; D1= 3.04 ± 0.08, D2 = 3.04 ± 0.08, D5 = 2.97 ± 0.08 mm). The variance in body size was also lowest in bees reared on D4, significantly lower than bees reared on D1 (Table S4; Brown-Forsythe Test *p* = 0.002) or D5 *(p* = 0.026). As expected, there was a significant positive relationship between body mass and body size (Fig. 2C, Table S5; *X*^2^_1_ =12.01, *p* < 0.001), but diet treatment had no significant effect on the relationship between body mass and body size.

Body condition scores (body mass/body size) also differed between diet treatments (Fig. 2D, Table 2, Table S3; F_3,65_ = 3.354, *p* = 0.024). Bees reared on the high carbohydrate diet (D2), had a significantly lower body condition score on average (emm = 26.1 ± 0.80 mg/mm) than those reared on D1 (emm = 29.5 ± 0.82 mg/mm; estimate ± s.e. 3.40 ± 1.14 mg/mm, *p* = 0.004) or D5 (emm = 28.8 ± 0.80 mg/mm; estimate ± s.e. 2.70 ± 1.13 mg/mm, *p* = 0.020). As with body mass, there was also a significant difference in the variance of body condition scores between bees raised on D2 and D1 (Table S4; Brown-Forsythe Test *p* = 0.011) and D2 and D5 (Brown-Forsythe Test *p* = 0.008).

### The ratio of P:C in larval diets affects the scaling of resting metabolic rate with body mass

Across all diet treatments, RMR (μL CO_2_ h^-1^) scaled positively with body mass (Fig. 3A, Table 3), and bees reared on the diet which maximised survival (D2) had a significantly steeper slope compared to those reared on D1 (Table 4). Diet also had a significant effect on the scaling of mass-specific RMR (RMR/body mass; Fig. 3B, Table 4). Bees reared on D2, the diet which maximised survival, showed a positive relationship between body mass and mass-specific RMR, whereas bees reared on all other diets exhibited a negative relationship (Fig. 3B). The difference in scaling between body mass and mass-specific RMR in bees reared on D1 and D2 was significant (Table 4; estimate ± s.e. = −1.00 ± 0.47, df = 92.03, *p* = 0.035). The nature of the scaling relationship between adult body mass and RMR differed considerably according to larval nutrition, with bees reared on D2 diet exhibiting positive allometry, bees reared on D5 diet exhibiting isometry and bees reared on D1 diet exhibiting negative allometry (Table 3). Body size was not a significant predictor of RMR (Fig. 3C; Table S5; F_1,53.36_ = 2.37, *p* = 0.242), and while for most diet treatments there was a positive relationship between body condition and RMR, again this was not a significant predictor (Fig. 3D, Table S5; F1, 63.72 = 2.67, *p* = 0.107).

**Figure 3.**
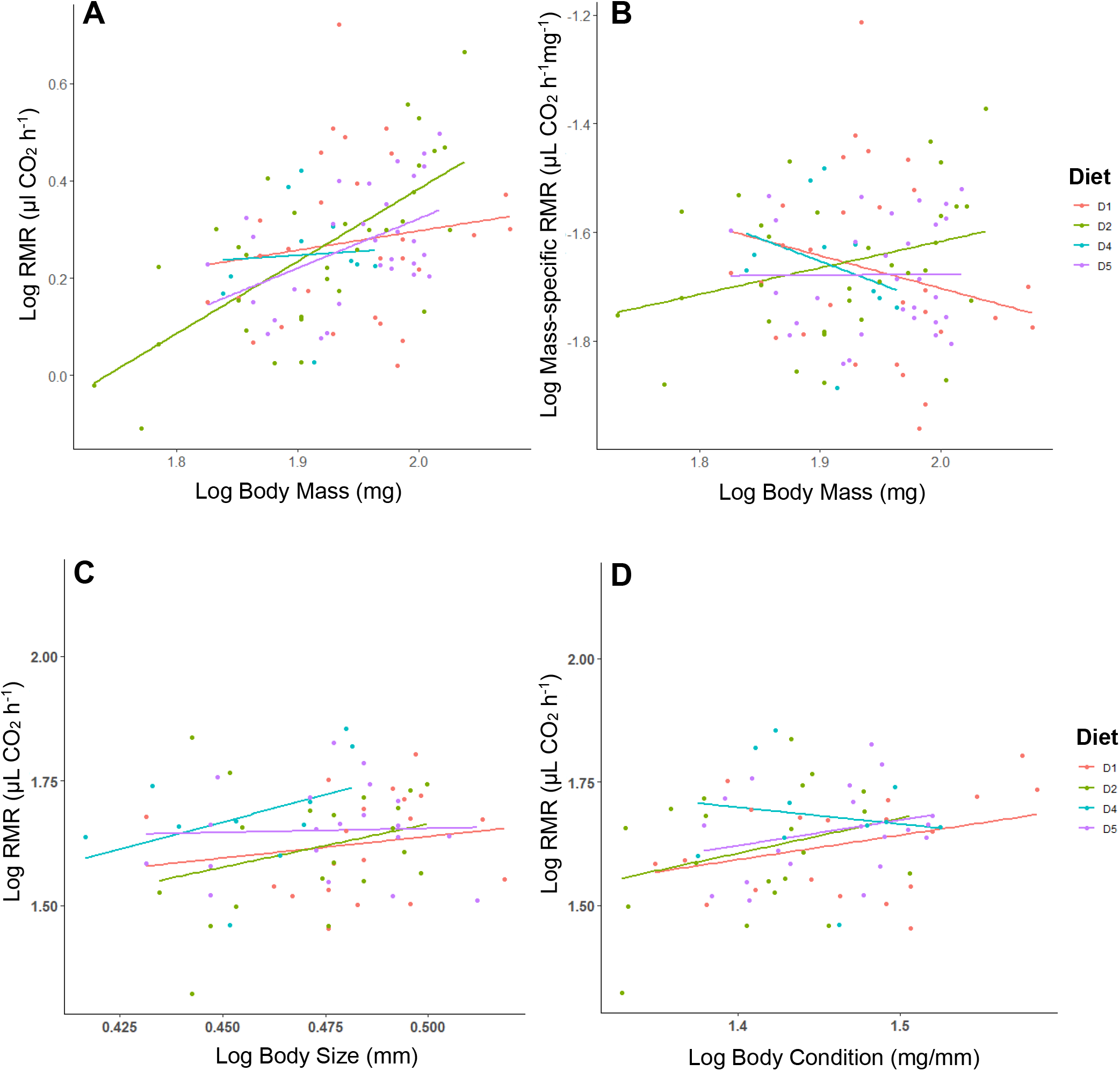
Scaling between CO_2_ production (RMR) and body mass (A), mass-specific RMR and body mass (B), RMR and body size (intertegular distance) (C) and RMR and body condition (body mass/body size) (D) for bees reared on larval diets differing in P:C ratio (D1 = 1:2.3; D2 =1:3.0; D3 = 1:1.5; D4 = 1:1.9; D5 = 1:2.9). The number of bees tested in each treatment is as follows: D1=28; D2=33; D3=0; D4=10; D5=30.

**Table 3.** Scaling relationship between body mass (mg) and RMR (μL CO2 h^-1^) for adult bees reared on different larval diets. Slopes and intercepts ± s.e. calculated *via* least-squares regression.

| diet | P:C | *n* | log slope ± s.e. | log intercept ± s.e. | regression equation |
| --- | --- | --- | --- | --- | --- |
| **D1** | 1:2.3 | 28 | 0.397 ± 0.52 | 2.155 ± 2.33 | $R^2 = 0.022$, $F_{1,26} = 0.585$, $p = 0.451$ |
| **D2** | 1:3.0 | 33 | 1.489 ± 0.28 | -2.674 ± 1.22 | $R^2 = 0.486$, $F_{1,31} = 29.27$, $p < 0.001$ |
| **D4** | 1:1.9 | 10 | 0.160 ± 0.94 | 3.168 ± 4.14 | $R^2 = 0.003$, $F_{1,8} = 0.029$, $p = 0.870$ |
| **D5** | 1:2.9 | 30 | 1.016 ± 0.36 | -0.636 ± 1.61 | $R^2 = 0.223$, $F_{1,28} = 8.035$, $p = 0.008$ |

**Table 4.** Effect of larval diet and body mass (mg) on RMR, and diet and body mass on mass-specific RMR. Models used were (log(RMR) ~ diet * log(body mass) + (1|grafting cohort)) and (log(mass- specific RMR)) ~ diet * log(body mass) + (1|grafting cohort)) respectively. The number of bees tested in each treatment is as follows: D1=28; D2=33; D3=0; D4=10; D5=30.

| | | estimate ± s.e. | d.f. | t-value | p-value | variance ± s.d. |
| --- | --- | --- | --- | --- | --- | --- |
| **RMR**<br>($\mu\text{L CO}_2 \text{ h}^{-1}$) | **fixed effects** | | | | | |
| | Intercept (D2) | -1.21 ± 1.27 | 92.16 | -0.95 | 0.343 | |
| | D1 | 4.36 ± 2.10 | 92.05 | 2.08 | **0.040** | |
| | D4 | 2.40 ± 4.63 | 92.46 | 0.52 | 0.610 | |
| | D5 | 4.02 ± 2.33 | 92.58 | 1.73 | 0.088 | |
| | (log)Body Mass | 1.17 ± 0.28 | 92.86 | 4.09 | **<0.001** | |
| | D1: (log)Body Mass | -1.00 ± 0.47 | 92.03 | -2.14 | **0.035** | |
| | D4: (log)Body Mass | -0.58 ± 1.05 | 92.43 | -0.55 | 0.582 | |
| | D5: (log)Body Mass | -0.93 ± 0.52 | 92.61 | -1.79 | 0.077 | |
|  | **random effects** |  |  |  |  |  |
| | grafting cohort | | | | | 0.04 ± 0.20 |
| | residual | | | | | 0.08 ± 0.29 |
| **mass-specific RMR**<br>($\mu\text{L CO}_2 \text{ mg}^{-1} \text{ h}^{-1}$) | **fixed effects** | | | | | |
| | Intercept (D2) | -1.21 ± 1.27 | 92.16 | -0.95 | 0.343 | |
| | D1 | 4.36 ± 2.10 | 92.05 | 2.08 | **0.040** | |
| | D4 | 2.40 ± 4.63 | 92.46 | 0.52 | 0.610 | |
| | D5 | 4.02 ± 2.33 | 92.58 | 1.73 | 0.088 | |
| | (log)Body Mass | 0.17 ± 0.28 | 92.86 | 0.58 | 0.563 | |
| | D1: (log)Body Mass | -1.00 ± 0.47 | 92.03 | -2.14 | **0.035** | |
| | D4: (log)Body Mass | -0.58 ± 1.05 | 92.43 | -0.55 | 0.582 | |
| | D5: (log)Body Mass | -0.93 ± 0.52 | 92.61 | -1.79 | 0.077 | |
|  | **random effects** |  |  |  |  |  |
| | grafting cohort | | | | | 0.04 ± 0.20 |
| | residual | | | | | 0.08 ± 0.29 |

## Discussion

Many organisms experience nutritionally sub-optimal diets during development, but very few studies have directly examined the impact of developmental diet quality on adult metabolism, particularly in insects. This question is of particular importance for bees, which as adult foragers face extremely high energetic demands, and as larvae experience a diet completely dependent on the provisioning choices of their mother and/or siblings, likely limiting their ability to self-regulate the intake of particular nutrients. Previous studies have shown that manipulating colony access to pollen results in reduced body size and life span in adult bees (Eishchen, Rothenbuhler and Kulinčević, 1982; Daly *et al*., 1995; Brodschneider and Crailsheim, 2010). Because the exact nutritional content of larval diets is manipulated at the colony level through the brood tending behaviour of nurse bees, larval nutrition is unknown in such studies. By using *in vitro* rearing methods, we were able to tightly control the macro-nutrient content of larval honeybee diets, demonstrating that the protein and carbohydrate content of the honeybee larval diet has a significant impact on larval development time, survival to adulthood, and adult body mass, size and condition. Using flow-through respirometry to measure whole-organism metabolism, we have shown for the first time that the protein and carbohydrate content of the larval diet of a holometabolous insect can impact the scaling relationship between adult body mass and RMR.

Larvae reared on a high carbohydrate diet had the highest survival to adulthood (D2, P:C 1:3), significantly higher than bees in all other treatment groups. Nearly all bees reared on the low carbohydrate diet failed to eclose (D3, P:C 1:1.5), and bees reared on the high protein diet (D4, P:C 1:1.9) also showed poor survival to adulthood. However, the absolute *amount* of protein and carbohydrate consumed over the course of development appears to be more important for survival than the ratio of macronutrients contained within the diet; though the low protein diet (D5, P:C 1:2.9) had a similar ratio of protein to carbohydrate as the high carbohydrate diet (D2), survival was significantly worse. Helm *et al*. (2017) also observed the highest survival in bees reared on a medium protein and high carbohydrate diet, and poor survival for bees reared on high protein diets, though survival was only recorded to the pupal stage. They concluded that there was an interaction between protein and carbohydrate on larval development, fitting with the idea of both the ratio and absolute amounts of protein and carbohydrate being important. Across all treatment groups, we observed most deaths occurring between pupation and adult eclosion, emphasising the importance of assessing survival to the adulthood. Pupation is a highly metabolically active period (Roeder and Behmer, 2014), suggesting that the impact of diet quality on the nutrient reserves available during this period may be the key to survival to adulthood.

The impact of high levels of dietary protein, both the absolute amount and relative content, upon survival has been demonstrated for bees as well as many other organisms (Lee *et al*., 2008; Dussutour and Simpson, 2009, 2012; Cook and Behmer, 2010; Pirk *et al*., 2010; Le Couteur *et al*., 2015; Solon-Biet *et al*., 2015). For example, the survival to adulthood, larval development, and size of solitary Megachilid bees is best on a high carbohydrate diet (Austin and Gilbert, 2018). The absolute quantity rather than the ratio of dietary macronutrients has also been shown to impact survival in soldier flies (Barragan-Fonseca *et al*., 2019). However, the mechanism underpinning the deleterious effect of consuming large volumes of protein on lifespan is poorly understood. It may be that digestion of large amounts of protein is very energetically costly (Westerterp, Wilson and Rolland, 1999; Halton and Hu, 2004), and produces toxic levels of nitrogen waste (Wright, 1995), though a recent study in adult ants has shown that even feeding free amino acids that require little digestion leads to a reduction in life span, leading the authors to suggest that excess amino acids may lead to an over-stimulation of the nutrient pathways that regulate lifespan (Arganda *et al*., 2017). Certain amino acids are seemingly more toxic than others, notably methionine, serine, threonine and phenylalanine, suggesting that the precise amino acid composition of an organisms’ diet may be important for survival and longevity.

Bees reared on the high carbohydrate diet, the best for survival, also took significantly longer to emerge as adults compared to bees in all other diet groups. This contrasts with previous studies in insects which have typically observed slower development on *lower* quality diets (Johnson, Wofford and Whitehand, 1992; Angell *et al*., 2020). For example, Rodrigues *et al*. (2015) found that when *Drosophila* larvae are given the choice between a diet that maximises survival, body size and fecundity, *versus* a diet that maximises development rate, they preferentially feed on the latter, potentially as a strategy to both avoid predation and maximise mating opportunities. Honeybees workers, in contrast, have a rather unique life history, developing inside a well-defended colony with much less risk of predation compared to other insects. Workers do not need to mate and reproduce, and there is little competition between individuals for resources which are provided for them by foragers and nurses. These factors may therefore reduce the pressure on honeybee larvae to develop quickly, particularly considering that rapid development is often thought to lead to earlier or faster senescence (Monaghan, Metcalfe and Torres, 2009). It remains to be determined how dietary macronutrients affect the development rates of the reproductive castes, queens and drones, that may face different pressures on developmental speed.

Diet also had a significant effect on emerging adult bees’ body mass, size and condition, which fits with previous studies linking the quality of pollen and nectar in larval diets to emergent adult bee size (Roulston and Cane, 2002; Burkle and Irwin, 2009). To our knowledge, however, this is the first study to demonstrate experimentally that the specific macro-nutrient composition of the larval diet affects body mass, size and condition in worker honeybees, which are typically considered to exhibit limited variation in body size compared to other bee species such as bumblebees (Goulson *et al*., 2002) or solitary bees. Perhaps unsurprisingly, bees reared on the worst diet for survival, the high protein diet, were the smallest and lightest on emergence. However, bees reared on this poor diet also had the narrowest range of body sizes, while those reared on the high carbohydrate diet had the best survival rate and the widest variation in body mass, suggesting that diets that increase survival also allow for a greater range of body sizes to emerge. Bees reared on the high carbohydrate diet had significantly lower body condition scores than bees reared on the diet containing a moderate amount of protein and carbohydrate. Diet-dependent variation in worker body size can have implications for both individual and colony functioning. Kerr & Hebling (1964) found that worker weight can affect the age with which worker honeybees make the transition from in-hive tasks to foraging, and in bumblebees and other bees, body size has been shown to correlate positively with foraging range (Greenleaf *et al*., 2007) and the weight of pollen and nectar loads that can be collected and transported back to the nest (Ramalho, Imperatriz-Fonseca and Giannini, 1998; Goulson *et al*., 2002; Kerr, Crone and Williams, 2019). Smaller bees have also been shown to be less effective at pollinating flowers (Jauker *et al*., 2012; Willmer and Finlayson, 2014). Thus, consuming inadequate amounts of macronutrients during development leads to both lower survival and body mass in adult worker bees, with potential consequences for the age structure and foraging efficiency of the colony, as well as wider ecological implications for the delivery of pollination.

Studies examining the impact of developmental diet on adult metabolism and metabolic scaling are rare, particularly in insects, and it is generally not agreed whether poor quality diets should lead to an increase or decrease in the RMR of emerging adults, given that this is likely to depend on the specific behavioural and/or physiological response(s) of an organism to an unbalanced diet (Burton *et al*., 2011). For example, organisms might be expected to reduce their metabolic rates in response to a low quality diet to minimise energetic expenditure (McNab, 1986). However, physiological adaptations to imbalanced diets, such as increasing gut length, may be metabolically costly (Yang and Joern, 1994). The few studies that have examined the impact of manipulating diet nutritional quality have generally found that nutritionally poor diets result in an elevation of the average RMR (14,33-35, but see 76). Typically these studies have considered only the short term impacts of diet on metabolism, during either adulthood or a single juvenile stage, and often do not report the impact of diet quality on the scaling of RMR with body mass or size. Here we demonstrated that body mass scales positively with RMR across all treatment groups, as is typical for insects (Niven and Scharlemann, 2005), and that larval diet has a long-term impact on metabolic scaling in adult bees. However, there were substantial differences in the slope of the scaling relationship between body mass and RMR depending on diet. Bees reared on the high carbohydrate diet (D2) showed positive allometry, with larger bees exhibiting higher RMR, which is very unusual [6-13]. In comparison, the RMR of bees reared on a diet containing a moderate amount of protein and carbohydrate (D1) showed isometry, whilst that of bees reared on the low protein diet (D5) exhibited a decelerating allometric relationship. Bees reared on the high carbohydrate diet (D2) also exhibited an unusual increase in mass-specific RMR with body mass, while bees reared on all other diets displayed a more typical decelerating or isometric relationship between mass-specific RMR and body mass. Neither body size (intertegular span) nor body condition scaled with RMR. This discrepancy with the finding for body mass is somewhat unexpected, given that we recorded RMR immediately following emergence, before additional feeding could strongly influence the bees’ mass. This is an important finding given that body size is often used as a proxy for body mass but may in fact scale quite differently with RMR. In contrast to our finding that diet quality affects the scaling of body mass and RMR, Karowe & Martin (1989) observed that while consumption of lower quality diets by larvae of the moth *Spodoptera eridania* led to an elevated RMR, the slopes of the positive scaling relationships between RMR and body mass were unaffected by diet treatment. However, in this study only protein quality was manipulated. Also metabolic rates were measured during the larval stage only, and in other organisms scaling relationships have been shown to change during ontogeny and could therefore be differentially affected by diet (Killen *et al*., 2007; Frappell, 2008). Consuming algal diets with unbalanced phosphorous:carbon ratios has been shown to change the scaling relationship between RMR and body mass in *Daphnia*, though this finding was based on a pooled data set across four closely related species (Jeyasingh, 2007). Therefore, our study is the first to demonstrate that the precise nature of an allometric scaling relationship can be altered by developmental diet within a single invertebrate species, significantly contributing to our understanding of the mechanistic basis of variation in the allometry of RMR (Vaca and White, 2010).

The use of CO_2_ production as a measure of RMR in the current study means that we were not able to detect changes in respiratory quotient arising from potential shifts in the substrate used for metabolism. For example, Clark *et al*. (2016) concluded that the lack of difference in RMR between winged and flightless morphs of the cricket *Gryllus firmus*, was due to a shift in metabolic substrate from carbohydrate to protein across diet treatments. Though we are unable to rule out this possibility, if such a shift has also occurred in the adult bees measured in our study, given that we tested bees on the day of emergence, prior to feeding, this would then suggest that substantial shifts in metabolic substrate use can also occur in response to differences in larval nutrition.

Here we considered only differences in protein and carbohydrate content of the diets, given the number of studies demonstrating that insect herbivores tightly control their intake of these two nutrients (Behmer, 2009). Royal jelly also contains around 5% lipids, on average, which are increasingly being recognised as an important component of larval nutrition, with bees appearing to regulate their intake of fats at both the level of the colony and individual foragers (Vaudo *et al*., 2016a; Vaudo *et al*., 2016b; Vaudo *et al*., 2020). Therefore, variation in the lipid content of the larval diet may also have had an impact on adult metabolism. Royal jelly also contains various micronutrients such as vitamins and sterols, which are important for hormone production and cannot be synthesised by bees themselves (Wright, Nicolson and Shafir, 2018). In one of the few other studies examining the impact of developmental diet on adult insect metabolism, Hill *et al*. (2020) observed changes in average RMR in stick insects reared from birth on leaves of three different plant species. Any effects on metabolic scaling were not reported. The macro-nutrient content of leaves from the three plant species did not show much variation, but the concentration and digestibility of micronutrients did. This suggests that in future studies, additional nutritional components other than the macro-nutrients protein and carbohydrate should also be considered in the context of dietary impacts on metabolism.

## Conclusions

There is increasing evidence that habitat fragmentation and farming intensification are reducing both the quantity and diversity of floral resources available for bees and other pollinators (Donkersley *et al*., 2017; Trinkl *et al*., 2020), which is of considerable concern given the global importance of insect pollination to ecosystem functioning and food security. Here we clearly demonstrate that the nutritional quality of larval diets impacts the metabolic functioning of adult worker bees, with diets more optimal for survival resulting in a higher metabolic rate per unit of body mass. As foraging bees already experience extremely high metabolic demands, differences in the quality of larval nutrition could impact the metabolic function which may negatively influence the foraging efficiency of workers. Subsequently this could impact on the build-up of pollen and nectar stores available for brood rearing and overwintering, with consequences for overall colony success.

## Acknowledgements

We thank Francis Ratnieks and Luciano Scandian for providing access to honeybee colonies and frames of larvae.

## Author Contributions

EN conceived the study, designed the methods, performed statistical analysis and wrote the paper. MR collected data in the laboratory, performed statistical analysis and commented on the paper. JN conceived the study and commented on the paper.

